# The cytoskeleton adaptor protein Sorbs1 controls the development of lymphatic and venous vessels in zebrafish

**DOI:** 10.1101/2020.08.20.259663

**Authors:** Alexandra Veloso, Anouk Bleuart, Tanguy Orban, Jonathan Bruyr, Pauline Cabochette, Raoul F.V. Germano, Alice Bernard, Benoit Vanhollebeke, Maud Martin, Franck Dequiedt

**Author notes:** Molecular Pathology Unit, Cancer Center and Regenerative Medicine, Harvard Stem Cell Institute, Massachusetts General Hospital Charlestown Navy Yard Campus, MA02129, USA. These authors contributed equally to this work (Alexandra Veloso, Anouk Bleuart). These authors also contributed equally to this work (Maud Martin, Franck Dequiedt).

## Abstract

Lymphangiogenesis, the formation of lymphatic vessels is tightly linked to the development of the venous vasculature, both at the cellular and molecular levels. Here, we identify a novel role for Sorbs1, the founding member of the SoHo family of cytoskeleton adaptor proteins, in vascular and lymphatic development in zebrafish. We show that Sorbs1 is required for secondary sprouting and emergence of several vascular structures specifically derived from the axial vein. Most notably, formation of the precursor parachordal lymphatic structures is affected in *sorbs1* mutant embryos, severely impacting the establishment of a proper trunk lymphatic network and leading to edema development. We show that Sorbs1 is probably not part of the Vegfc signaling, but instead might interacts with the BMP pathways. Mechanistically, we show that Sorbs1 controls FAK/Src signaling to impact on Rac1 and RhoA GTPases-regulated cytoskeleton processes. Inactivation of Sorbs1 altered cell-extracellular matrix (ECM) contact rearrangement and cytoskeleton dynamics, leading to specific defects in endothelial cell migratory and adhesive properties. Our data thus establish Sorbs1 as an important regulator of lymphangiogenesis distinct from the Vegfc signaling axis, increasing our understanding of context-specific vascular and lymphatic development.

## Introduction

The adult circulatory system encompasses the blood and lymphatic vasculatures. Intimate connections link these networks at the developmental, anatomical and functional level. In the embryo, the blood vasculature develops through a sequence of events that have been extensively characterized in the past decades^1^. Our understanding of lymphangiogenesis, the formation of lymphatic vessels, lags far behind that of angiogenesis, the formation of new blood vessels. Because of its critical role in tissue homeostasis and immune surveillance lymphangiogenesis has gained a lot of attention in recent years, leading to the identification of an increasing yet limited number of molecular players. Beyond these advances, recent reports also pointed to the protective and reparative therapeutic potential of enhancing lymphangiogenesis in several pathological contexts such as myocardial infarction^2–4^, glioblastoma^5^ and renal dysfunction^6^.

As for vascular development, the zebrafish model has significantly contributed to expend our understanding of lymphatic system formation and biology. More specifically, the stereotypical formation of the trunk lymphatic network has been extensively used to decipher the driving principles of lymphangiogenesis. Trunk lymphatic endothelial cell (LEC) precursors arise from transdifferentiation of venous ECs located within the posterior cardinal vein (PCV). Specification of LECs is triggered by the expression of the transcription factor Prox1a and occurs before their effective egression from the vein in a process involving asymmetric division^7–9^ and regulated by transcriptional ^10–14^ and post-transcriptional progams^15^. At 32-34 hours post-fertilization (hpf), dorsal sprouting of these lymphatic-fated cells contributes to the transient development of a longitudinal string of parachordal lymphangioblasts (PLs) 20 h later. At around 60 hpf, parachordal LECs start to migrate ventrally and dorsally to form the major trunk lymphatic network consisting of the thoracic duct (TD), the intersegmental lymphatic vessels (ISLVs) and the dorsal longitudinal lymphatic vessels (DLLV)^16^.

Illustrating the striking plasticity of endothelial cells, not all the vascular sprouts emerging from the PCV and migrating dorsally alongside the artery-derived primary intersegmental vessels (aISVs) participate in building the lymphatic vessels. Approximately half of them, almost undistinguishable except for their reduced Prox1a expression, will connect and anastomose to the proximal region of aISVs to form venous ISVs (vISVs)^16–18^. Adding to the behavioral heterogeneity and specialization of the venous ECs from the PCV, ventral angiogenic sprouting also occurs from the posterior and anterior parts of the axial vein at different time points. ECs in the caudal region sprout from the floor of the caudal vein (CV) at around 27 hpf and migrate towards the ventral side of the embryo to form the caudal vascular plexus (CVP), a distinctive fenestrated network of vessels^19^. More rostrally, formation of the subintestinal venous plexus (SIVP), which will eventually provide blood supply to the digestive tract starts at around 30 hpf with a process of ventral migration leading to the formation at 3 days post-fertilization (dpf) of a basket-like plexus composed of vertical interconnecting vessels (ICVs) that drain into a transversal subintestinal vein (SIV)^20,21^.

Originating from the same parental vessel, the process of trunk lymphangiogenesis is tightly intermingled with PCV-derived venous angiogenesis and especially with ISV secondary sprouting that occurs concomitantly. Studies of mutants isolated from forward genetic screens or associated with human diseases led to the establishment of the Vegfc/Flt4 axis as the central pathway for lymphangiogenesis^22–25^. Accordingly, the currently growing list of lymphangiogenesis regulators almost exclusively relates to molecules involved in Vegfc/Flt4 signaling, some of them acting directly upstream such as CCBE1^26–29^ or the transcription factor HHEX^11^ or downstream like the transcription factors Mafba^7^ and Yap1^30^. Acting through multiple intracellular events, including the activation of the common effector of Vegf receptors Erk^8^, Vegfc signaling has been shown to control several aspects of lymphangiogenesis including LEC differentiation through Prox1 expression, proliferation and migration. Whereas these signaling cues seem to act indistinctly on lympho-venous sprouting, with the majority of effectors impacting on both lymphatic vessel and vISV formation^22,27,31,32^, ventral angiogenesis of the caudal vascular and subintestinal plexus specifically relies on BMP signaling^19,21^.

While our understanding of angiogenic cues that drive formation of specific vascular beds is only emerging, even less is known about the intracellular components that define specific endothelial cell behavior during establishment of highly conserved organ-specific vascular patterns. Sorbs1 (Cbl associated protein CAP/ponsin) belongs to the SoHo family of adaptor proteins that includes two other members, Sorbs2 (Arg-binding protein 2, ArgBP2) and Sorbs3 (Vinexin). Early following their discovery, SoHo proteins were shown to localize to various actin-based structures, including z-discs, stress fibers, cell-ECM and cell-cell adhesions^33–39^. Sorbs1 interactions with several structural and signaling cytoskeletal components, such as vinculin and paxillin, strengthened the idea that it might function as an adaptor protein coordinating multiple signaling complexes regulating the actin cytoskeleton^40,41^. In agreement with these observations, *in vitro* studies showed that Sorbs1, along with the other family members are important regulators of actin-dependent processes, such as migration, adhesion and mechano-transduction^39,42,43^. Such cytoskeleton-based processes have been shown to be essential to support and control the morphogenic events that endothelial cells have to go through during blood and lymphatic vessel formation^44^.

Little is known about the *in vivo* biological functions of SoHo proteins. Here, we report that Sorbs1 has unsuspected roles in zebrafish developmental angiogenesis and lymphangiogenesis. Using a combination of *in vivo* and *in vitro* approaches, we demonstrate that Sorbs1 controls endothelial cell adhesion signaling through modulation of specific RhoGTPases activities and consequently participates in the formation of specific venous and lymphatic structures originating from the main axial vein, both dorsally and ventrally. Surprisingly, despite its major impact on trunk lymphatic structures, Sorbs1 is not involved in Vegfc pathway but appears to participate in BMP signaling.

## Results

### Sorbs1 genetic depletion is associated with pericardial edema formation

To investigate the function of Sorbs1 *in vivo*, we took advantage of the zebrafish model. We performed phylogenetic analysis using results from BLAST homology searches against NCBI and Ensembl databases and identified SoHo family orthologs in zebrafish. A single Sorbs1 ortholog (ENSDARG00000103435), two Sorbs2 orthologs, *sorbs2a* (ENSDARG00000003046) and *sorbs2b* (ENSDARG00000061603) and a single Sorbs3 ortholog (ENSDARG00000037476) were identified (Supplementary Figure S1A). To assess the role of Sorbs1 in zebrafish development, we used the CRISPR/Cas9 system to generate a *sorbs1* mutant allele. We selected F1 heterozygous carriers with a 14 base pair (bp) frame-shift deletion at codon 178, in the region of the *sorbs1* gene coding for the SoHo domain. This mutation is predicted to generate a premature stop codon at codon 182 (Supplementary Figure S1 B) and *sorbs1* homozygous mutants (referred to as *sorbs1^-/-^*) from heterozygous in-crosses express no detectable Sorbs1 protein (Supplementary Figure S1C). *Sorbs1* mutants exhibited no gross morphological abnormalities. Nevertheless, we observed that a large proportion of the *sorbs1^-/-^* larvae exhibited large edemas around the heart and the intestinal tract, which were first detectable at 2 dpf and were clearly visible at 5 dpf (Figure 1A and 1B). We suspected that theses edemas could be indicative of vascular and/or lymphatic defects, although *sorbs1* mutants displayed normal heart rates (Supplementary Figure S1D) and overall circulation. The presence of edema strongly impacted on the viability of the embryos with about 40% of edema-developing embryos dying within 10 dpf (Figure 1C). As a complementary approach, we also used a splice-blocking antisense morpholino (*sorbs1* MO) targeting the exon 3/intron 3 boundary of *sorbs1.* This morpholino efficiently prevented splicing of intron 3 and reduced Sorbs1 protein levels when injected at 5 ng/embryo (Supplementary Figure S1E,F). Similarly to *sorbs1* mutants, the vast majority of morphant embryos exhibited edemas, a defect that was rescued by injection of RNA coding for the human Sorbs1 ortholog (Supplementary Figure S1G). Based on these observations, we examined the expression of Sorbs1 in endothelial cells. Sorbs1 expression was detected *in vivo* in blood vessel endothelium by immunohistochemical analysis of various human tissues (Supplementary Figure 1H, black arrows). In addition, Sorbs1 protein was detected in various cultured human ECs, with the highest levels being observed in venous and lymphatic ECs (Supplementary Figure 1I). In zebrafish, whole mount *in situ* hybridization revealeded a ubiquitous expression of *sorbs1* throughout development (Supplementary Figure 1J). To validate *sorbs1* expression in the zebrafish vascular endothelium, we used the *Tg(fli1a:eGFP)y1* transgenic line, in which lymphatic, arterial, and venous ECs express green fluorescent protein (GFP), and sorted ECs (*i.e.*, GFP-positive cells) and non-ECs (*i.e.*, GFP-negative cells) by flow cytometry. Quantitative PCR analysis revealed that expression of *sorbs1* was significantly higher in ECs, as compared to non-ECs (Figure 1E). The expression of *sorbs1* in ECs was maximal at around 48 hpf, when active lympho-venous sprouting is occurring (Figure 1F).

**Figure 1:**
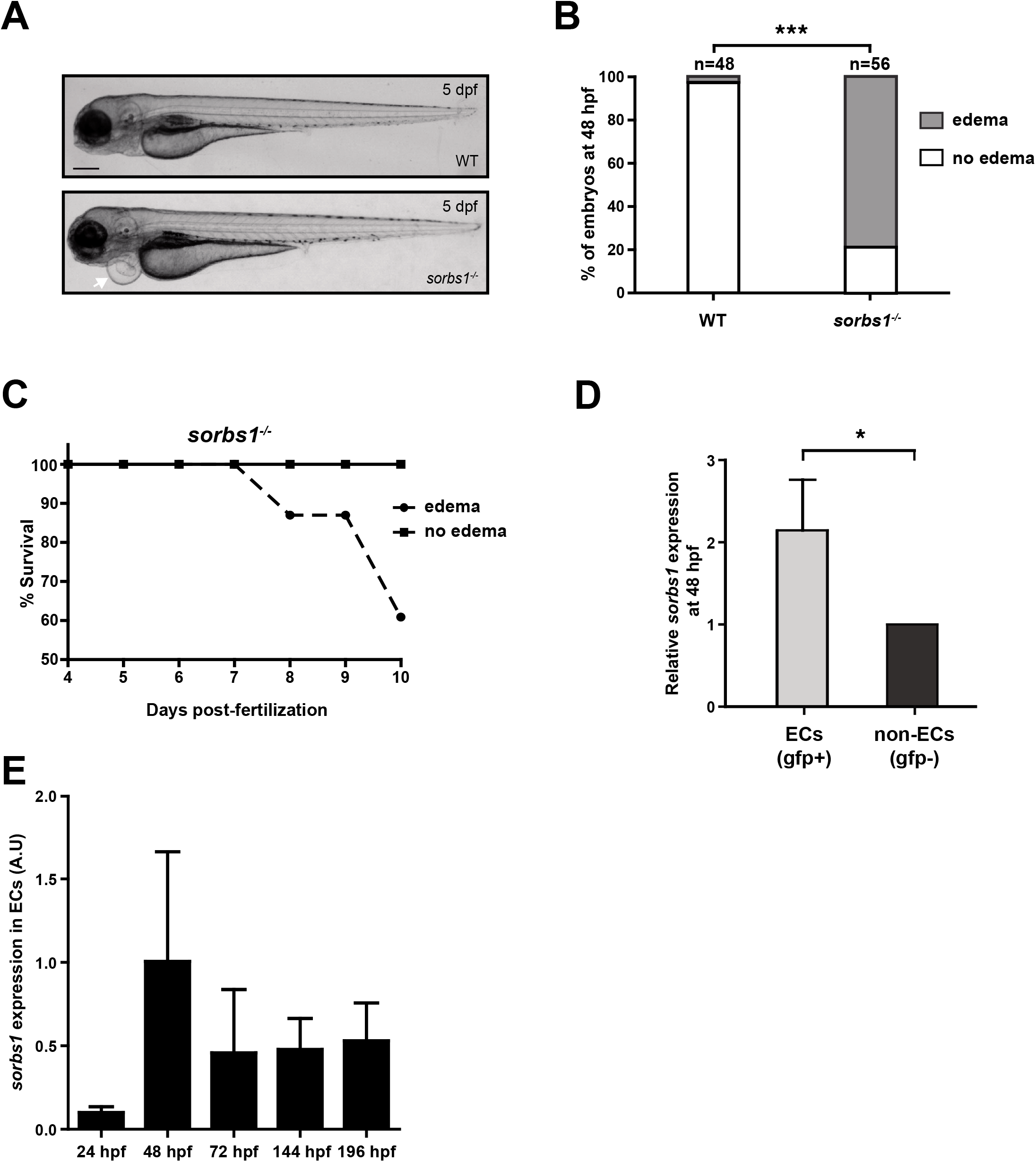
*Sorbs1* expression is enriched in the endothelium and its knockout in zebrafish results in cardiac edemas. (A-B) Transmitted light images of live wild-type (WT) and sorbs1 mutants (*sorbs1^-/-^*) zebrafish embryos at 5 days post-fertilization (dpf) (A) and quantification of the percentage of embryos displaying edemas (B). (n=number of embryos; *** *P<0.001;* Fischer exact test). The arrow indicates an example of edema observed in *sorbs1^-/-^* embryo. Scale bar represents 250 μm. (C) Quantification of the percentage of survival for *sorbs1^-/-^* embryos presenting or not edemas from 4 to 10 dpf. (n=24 and n=23, respectively). (D-E) RT-qPCR analysis of *sorbs1* expression relative to Elfa at 48hpf in endothelial cells (ECs, GFP+) vs non-endothelial cells (non-ECs, GFP-) sorted from WT *Tg(fli1a:eGFP)y1* embryos by FACS (Fluorescence Activated Cell Sorting) technology (D). Relative endothelial expression of Sorbs1 was quantified at different time points of embryonic development (E). (*P<0.05, Unpaired t-test).

### Sorbs1 is important for lymphangiogenesis in zebrafish

To explore if the presence of edema in *sorbs1^-/-^* larvae could relate to lymphangiogenesis deficiency, we performed microscopic observation of the vasculature of *sorbs1* mutants in the *Tg(fli1a:eGFP)y1* transgenic background. Whereas, mutants showed normal morphogenesis, patterning and lumenization of the cranial and trunk primary vasculature, development of lymphangiogenic structures was severely affected in the absence of *sorbs1* (Figure 2A, Supplementary Figure S2A,B). The formation of the PLs at the horizontal trunk septum was strongly impaired: quantification analysis at 54 hpf, confirmed that the proportion of somite segments with detectable PLs was significantly reduced in *sorbs1^-/-^* embryos, with PLs being totally absent in approximately one third of the embryos (Figure 2B). Similar defects were also detected in *sorbs1* morphants (Supplementary Figure S2C,D). At approximately 60 hpf, PLs migrate ventrally from the horizontal myoseptum to form the TD (3-6 dpf), the major lymphatic trunk vessel situated between the DA and PCV. Because *sorbs1* mutants had a lower number of PLs, we reasoned that they might also exhibit defects in TD formation. To test this, we measured the length of visible TD portions in 10 somites at 4 and 6 dpf, and expressed it as a percentage of the total length of this trunk segment (Figure 2C,D)^45^. In control larvae, the observed length of the TD at 4 dpf represented approximately 49% of the trunk total length, a proportion that increased up to 60% at 6 dpf (Figure 2D). Formation of the TD was greatly impaired in *sorbs1^-/-^* larvae, as TD length corresponded to only 19% and 26% of the 10-somite length at 4 and 6 dpf, respectively. In a large proportion of *sorbs1^-/-^* larvae (41%, 24/58) the TD was totally absent at 4 dpf while only 14% (8/57) of control embryos had no detectable TD. As expected, the most affected *sorbs1^-/-^* embryos (*i.e.*, embryos with less than 20% of visible TD) had reduced life span (Figure 2E). The defects in PL and TD formation strongly suggest that *sorbs1* is important for early lymphatic development, its absence culminating in edema formation and higher embryonic mortality. Importantly, in agreement with the idea of an endothelial function for Sorbs1, PL formation defects in *sorbs1^-/-^* mutants were rescued through endothelial specific ectopic expression of human Sorbs1 (Figure 2F,G).

**Figure 2:**
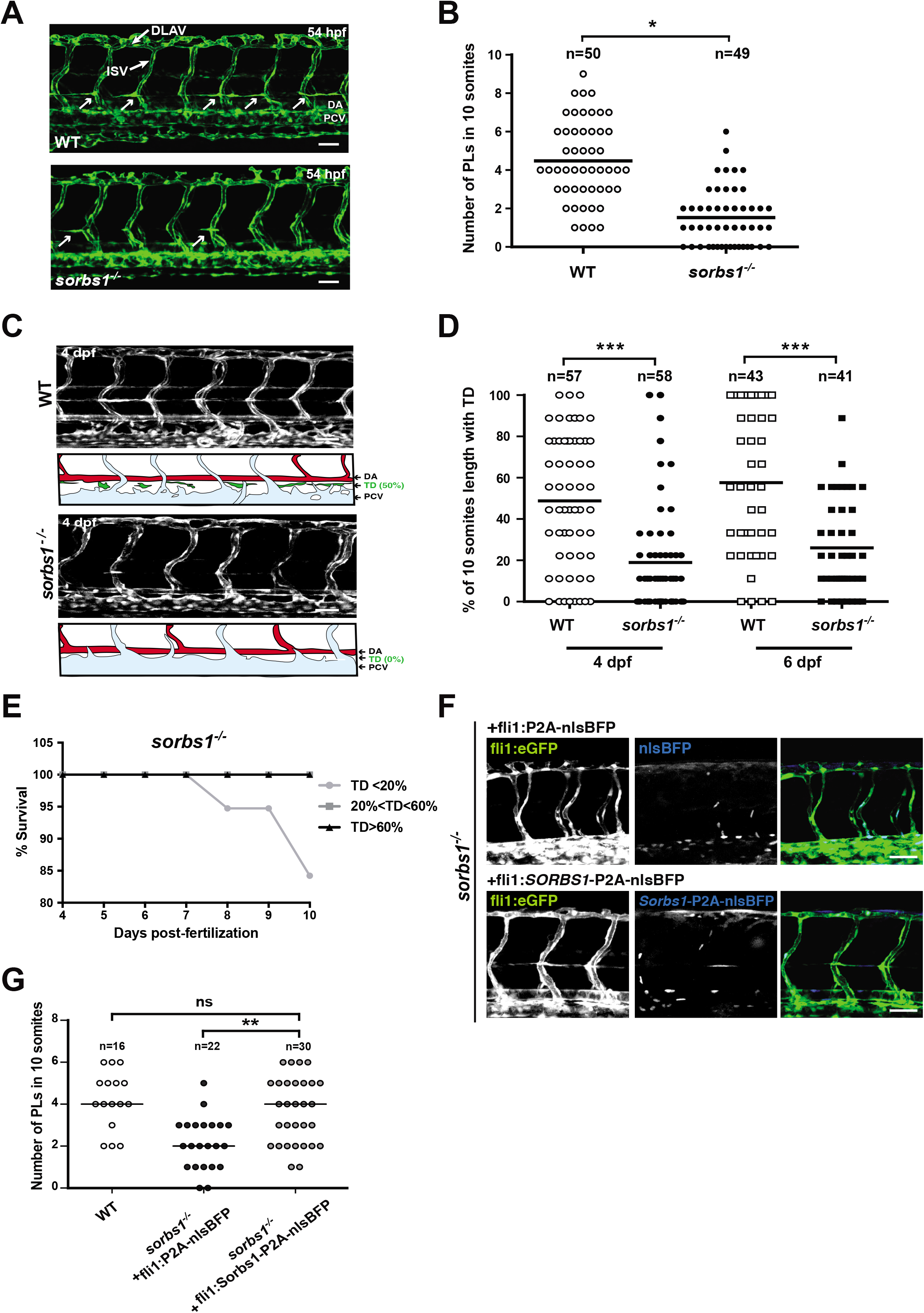
Sorbs1 is necessary for trunk lymphangiogenesis *in vivo*. (A-B) Confocal microscopy analysis of the trunk vasculature in WT and *sorbs1^-/-^ Tg(fli1a:eGFP)y1* embryos at 54 hpf (A) used to quantify the number of paracordal lymphangioblasts (PLs) (white arrows in A) over 10 somite segments (B). Dorsal aorta (DA), post cardinal vein (PCV), intersegmental vessels (ISV) or dorsal longitudinal anastomotic vessels (DLAV). (n=number of embryos; *P<0.05; Mann–Whitney *U*-test). Scale bars represent 50 μm. (C) Z-maximum projections of confocal images of the trunk vasculature from 4 dpf *Tg(fli1a:eGFP)y1* WT or *sorbs1* knock-out embryos. Schematic representations of arterial (red), venous (light blue) and lymphatic (green) vessels are showed below the confocal pictures. Dorsal aorta (DA), Posterior Cardinal Vein (PCV), Thoracic duct (TD). Scale bars represent 50 μm. (D) Quantification of the thoracic duct (TD) extent over 10 segments at 4 and 6 dpf in WT and *sorbs1^-/-^* embryos. (n=number of embryos; ***P<0.001; Mann–Whitney *U*-test). (E) Analysis of *sorbs1^-/-^* embryo survival over time in relation to their TD defects. (F-G) Z-maximum projections of confocal images of the trunk vasculature of 54 hpf *Tg(fli1a:eGFP)y1 sorbs1* knock-out embryos expressing transgenic endothelial constructs coding for human Sorbs1 or not (F) and quantification of the number of PLs in the indicated condition (G). BFP is used as transgenesis marker. (n=number of embryos, ns=non-significant; *\*\*P<0.01;* Mann–Whitney *U*-test). Scale bars represent 50 μm.

### Sorbs1 function in lymphangiogenesis is independent of Vegfc

Almost all currently known genetic regulators of zebrafish trunk lymphangiogenesis, act through the Vegfc signaling pathway, the major regulator of lymphangiogenesis^46,47^. Vegfc induces expression of Prox1a in a subset of ECs in the PCV during lymphatic specification, triggering their sprouting, migration and proliferation to form the lymphatic trunk vessel network. qPCR analysis of *prox1a* expression in ECs showed no significant difference between wild-type and *sorbs1* mutants at 48 hpf (Figure 3A). Moreover, live imaging of *TgBAC(prox1a:KalTA4-4xUAS-ADV.E1b:TagRFP)^nim5^* line confirmed the presence of Prox1a-positive endothelial cells in the PCV of *sorbs1^-/-^* mutants (Figure 3B). However in *sorbs1^-/-^* mutants, Prox1a-positive cells failed to sprout out of the axial vein, indicating that *sorbs1* is dispensable for lymphatic specification but seems to be required for subsequent migration of LECs (Figure 3B). To directly test a link between Sorbs1 and Vegfc signaling we analyzed the lymphatic network of double *sorbs1/vegfc* heterozygous zebrafish embryos as haploinsufficiency has been demonstrated for Vegfc in this functon^29^. We found no evidence of genetic interaction between these two genes (Figure 3C). In line with these findings, injection of a Vegfc-coding RNA in *sorbs1* mutant embryos increased formation of PLs and TD similarly to control embryos, indicating that *sorbs1^-/-^* ECs are not affected in their potential to respond to ectopically produced Vegfc (Figure 3D, Supplementary Figure S3A).

**Figure 3:**
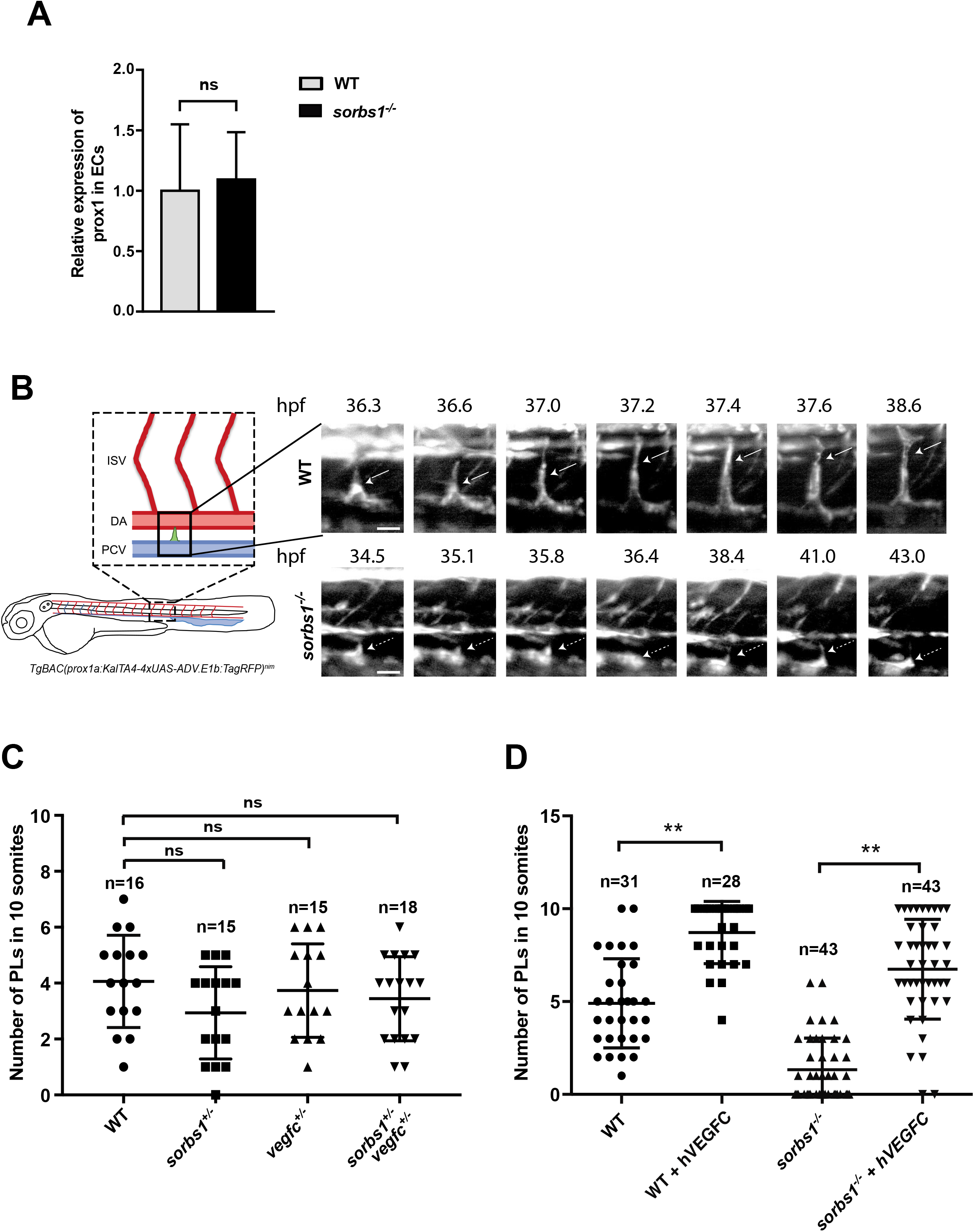
Sorbs1 functions independently of Vegfc signaling during *in vivo* lymphangiogenesis. (A) RT-qPCR analysis of *prox1a* relative expression at 48hpf in endothelial cells (ECs) sorted from WT and *sorbs1^-/-^ Tg(fli1a:eGFP)y1* embryos by FACS (Fluorescence Activated Cell Sorting) technology. (ns= non-significant, Unpaired t-test). Results are means from 5 experiments. (B) Frames (Z-maximum projections) from time-lapse lightsheet imaging of prox1 expressing ECs sprouting from the PCV in WT and *sorbs1^-/-^ TgBAC(prox1a:KalTA4-4xUAS-ADV.E1b:TagRFP)^nim5^* embryos. White arrows point to Prox1-positive ECs that sprouted to form PLs in WT but failed to migrate dorsally in *sorbs1^-/-^.* Scale bars represent 25 μm. (C) Quantification of PL extent within the trunk region of 54 hpf *Tg(fli1a:eGFP)y1* embryos from the indicated genotype resulting from the incross of *sorbs1^+/-^* with vegf^+/-^ embryos (n= number of embryos; ns=non-significant; Mann–Whitney *U*-test). (D) Quantification of the trunk PL (54 hpf) in WT or *sorbs1^-/-^ Tg(fli1a:eGFP)y1* embryos injected or not with human Vegfc. (n= number of embryos; **P<0.01; Mann–Whitney *U*-test).

### Lack of Sorbs1 impairs secondary sprouting from the PCV

In parallel to migration of Prox1a-specified cells to form PLs, venous Prox1a-negative ECs sprout from the PCV to connect to arterial intersegmental vessels (aISVs). To evaluate the role of Sorbs1 in this process, we counted the nascent secondary sprouts emerging from the PCV at 34 hpf, *i.e.* sprouts not yet fused to a primary ISV or stabilized to form PL (arrow in Figure 4A). Compared to control, the number of sprouts was significantly reduced in *sorbs1^-/-^ Tg(fli1a:eGFP)y1* embryos (n=77, P<0.01) (Figure 4B). The vast majority of embryos (68.8%) had no visible secondary sprouts and in the remainder fraction, only 15.6% of *sorbs1^-/-^* embryos had one secondary sprout, whereas 15.6% had 2 or 3. These defects in secondary sprouting were confirmed by looking at *sorbs1* morphants (Supplementary Figure S4A). Because approximately half of the secondary sprouts gives rise to PLs, while the other half connects to the primary ISV network and establish vISVs, the defective PCV secondary sprouting in the absence of *sorbs1* could explain the reduced number of PLs. To assess if it also affected migration of the venous sprouts that will connect to and remodel ISVs, we counted the number of vISVs, *i.e.* ISVs connected to the PCV over a 10-somite region (Figure 4C). In wild-type embryos, slightly less than half of the ISVs were connected to the PCV and thus scored as of venous identity. By contrast, the proportion of vISVs was significantly lower (35.5%) in *sorbs1* mutant embryos (Figure 4D). A similar reduction in vISVs was also observed in *sorbs1* morphants (Supplementary Figure S4B). These observations suggest that *sorbs1* knockout/knockdown affects the secondary wave of migrating ECs from the PCV, which is associated with both lymphatic and vISV network formation. In contrast, *sorbs1* is dispensable for primary sprouting from the DA.

**Figure 4:**
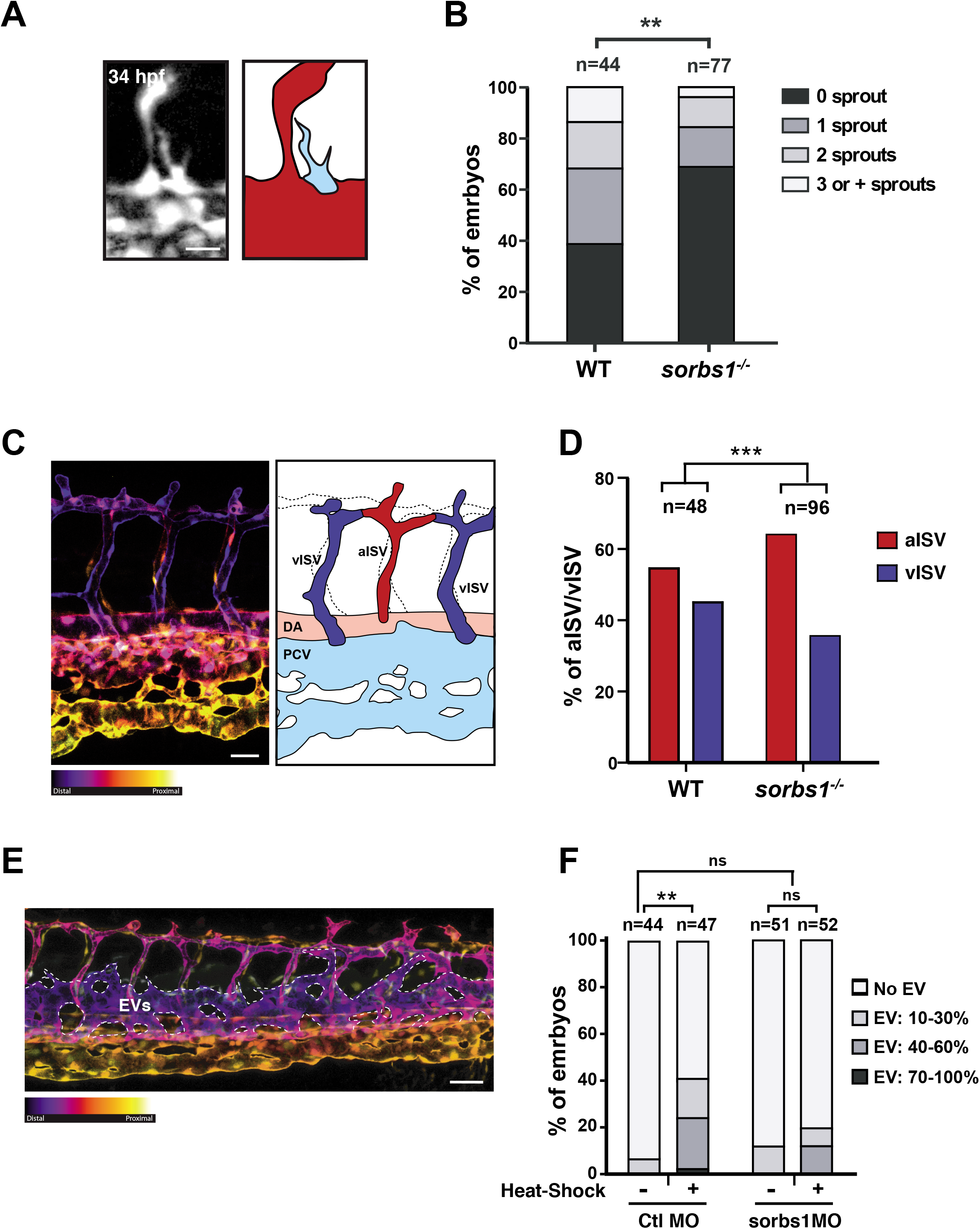
Sorbs1 depletion results in defects in secondary sprouting from the PCV. (A) Confocal image (top) and schematic drawing (bottom) of secondary sprouts (arrow) emerging from the PCV in *Tg(fli1a:eGFP)y1* embryos at 34 hpf. The DA and the primary ISVs are in red and the PCV is in blue. Scale bars represent 25 μm. (B) Quantification of secondary sprouts visible at 34 hpf in WT and *sorbs1* mutants (sorbs1^-/-^). (n=number of embryos, **P<0.01; Mann–Whitney *U*-test). (C-D) Color-coded Z-maximum projections of confocal images of the trunk regions of *Tg(fli1a:eGFP)y1* WT embryos and its schematic representation (C). The depth-associated color scale (warm colors = deep, cold colors = surface) allowed distinction between vISVs and aISVs to quantify their proportion at 48 hpf in a 10 somites trunk region of WT or *sorbs1^-/-^* embryos. (n= number of embryos, *** *P< 0.001;* χ2 with Yates correction). Dorsal aorta (DA), light post cardinal vein (PCV), arterial Intersegmental vessel (aISV) venous intersegmental vessel (vISV). Scale bars represent 50 μm. (E) Confocal image representation as described in C of 54hpf *Tg(hsp70l:bmp2b; fli1a:eGFP*) heat-shocked embryos that were heat-shocked at 26 hpf used to illustrate formation of ectopic vessels (EVs, indicated with dotted lines) (A). Scale bars represent 50 μm. (F) Quantification of ectopic vessel (EV) growing from the PCV at 28 hpf in *Tg(hsp70l:bmp2b; fli1a:eGFP*) embryos injected with Ctl or *sorbs1* Mo before (-) or after (+) a heat-shock treatment at 26hpf. (n=number of embryos, ** *P<0.01*, ns=non-significant, χ2 pairwise proportion test with Holm correction).

Along with the angiogenic dorsal sprouts, additional vascular structures are established from the PCV (Supplementary Figure S4C). Starting at 25 hpf, venous angiogenic sprouts emerge in the caudal region of the PCV and migrate ventrally through active angiogenesis to form the primordial caudal vein plexus (CVP), a complex network of vessels. During this process, ECs from the caudal vein extend protrusions towards the ventral region of the trunk to migrate and connect with each other to form the CVP at 48 hpf. We observed that while forming, the CVP from *sorbs1* mutant and knock-down embryos produced fewer ventral sprouts (Supplementary Figure S4D,E). Sprouting angiogenesis also occurs in the anterior region of the PCV leading to the formation of the SIVP. Several studies have extensively described the development of the SIVP and demonstrated that it forms from cells originating from the ventral side of the PCV, at around 30 hpf^20^. These ECs collectively engage in a process of ventral migration and give rise at 3 dpf to a left and right basket-like plexus composed of vertical interconnecting vessels (ICVs) that drain into a transversal subintestinal vein (SIV) (Supplementary Figure S4F). Formation of the subintestinal venous plexus was affected in the absence of *sorbs1. Sorbs1^-/-^* embryos showed abnormal SIV morphology, with irregular branching and in severe cases, absence of the surrounding SIV (Supplementary Figure S4F). In sum, phenotypic characterization of *sorbs1* morphants and mutants revealed phenotypes linked to defects in the development of every major angiogenic structure that originates from the PCV. Interestingly, some of these processes rely on the Bone Morphogenetic Protein (BMP) pathway. More specifically, BMP signaling promotes ventral venous sprouting during CVP development and collective EC migration during SIV ventral expansion^21^. Its role during dorsal secondary sprouting is less clear. To examine the role of Sorbs1 in BMP-induced venous angiogenesis, we used the *Tg(hsp70l:bmp2b*) line, in which ectopic endothelial sprouting can be specifically induced from the PCV by heat-shock treatment (Figure 4E). When double transgenic *Tg(hsp70l:bmp2b; fli1a:eGFP*) embryos were heat-shocked at 39°C for 30 min at 26 hpf (*i.e.*, at the onset of PCV secondary sprouting), 40% showed ectopic vessels (EVs). Sorbs1 knockdown significantly reduced Bmp-induced sprouting from the PCV, since less than 20% of *sorbs1* morphants displayed EVs after heat-shock (Figure 4F). In these embryos, EVs were also visible in a smaller proportion of somite segments, demonstrating that *sorbs1* is implicated in the venous EC response downstream or acting in parallel to BMP.

### Sorbs1 controls EC adhesion through regulation of small RhoGTPases

In order to understand the cellular and molecular mechanisms underlying Sorbs1 function during venous sprouting, we generated primary venous ECs deficient for Sorbs1 using small interfering RNA (siRNA) that efficiently and specifically suppresses the expression of Sorbs1, without affecting the viability or proliferation of ECs (Supplementary Figure S5A-D). In agreement with the observed impairment in EC migration from the PCV *in vivo*, downregulation of Sorbs1 correlated with a significant decrease in EC *in vitro* migratory capacities (Supplementary Figure S5E,F).

Members of the SoHo family are thought to function by interacting with and coordinating the activity of actin cytoskeleton regulators, including RhoGTPases^39,48–51^. During zebrafish CVP formation, BMP has been shown to affect EC migration by promoting endothelial filopodia extension via activation of Cdc42^71^. We thus assessed the activity of Cdc42 by performing Rho GTPase activity assays in control and Sorbs1-depleted ECs. Levels of active Cdc42 were similar in the presence or absence of Sorbs1 (Figure 5A). In contrast, when looking at the other Rho GTPase members, we found that knockdown (KD) of Sorbs1 correlated with a significant up-regulation in RhoA and a marked decrease in Rac1 activities. Reduction in Rac1 activity was associated with reduced phosphorylation of Rac1 effector kinases PAK2 and PAK4 (Supplementary Figure S5G). Activation of RhoA was confirmed by looking at actin polymerization at the lamellipodia of spreading Sorbs1-KD cells. Indeed, cells depleted for Sorbs1 exhibited a denser network of actin bundles at the cell periphery and treatment with the C3 Transferase Rho inhibitor prevented appearance of peripheral F-actin in Sorbs1-KD cells, confirming the causative role of RhoA (Figure 5B,C). To get more insight into the cellular function of Sorbs1, we checked its subcellular localization in ECs and found that it localizes at cell-ECM adhesions (Supplementary Figure S5H). The formation and maturation of integrin adhesions at the leading edge of migrating cells is controlled by a precise spatio-temporal balance between the activities of Rac1 and RhoA GTPases^52^. Rac1 promotes the formation of new adhesions in regions of membrane protrusions, but also regulates adhesion turnover through downstream effectors such as PAKs and local inhibition of RhoA^53^. In contrast, RhoA activation is associated with actomyosin-dependent stabilization and maturation of adhesions^52,54^. We examined the possibility that Sorbs1 might control EC adhesion dynamics by modulating the activity of Rac1 and RhoA. Inactivation of Sorbs1 resulted in alterations in the pattern of EC-ECM adhesions (Figure 5D). Cell-ECM adhesions found at membrane protrusions are usually divided into two types, depending on their maturation stage. The first adhesions to appear are nascent adhesions (NA) and focal complexes (Fx), which are small dot-like structures characterized by their high content in tyrosine-phosphorylated signaling molecules, such as phospho-Paxillin^55^. Few of them will elongate centripetally and mature into larger (area > 1 μm^2^) focal adhesions (FAs), in a process relying on actin filaments^54^. Compared to control siRNA-treated ECs, *Sorbs1* KD cells had a higher proportion of large FAs, which were localized more centripetally (Figure 5D,E). In contrast, the number of small phospho-Paxillin positive adhesions was reduced at the periphery of Sorbs1-deficient cells (Supplementary Figure S5I,J). Importantly, the excessive accumulation of stable FAs was correlated with a significant increase in cell adhesion onto fibronectin, providing a potential explanation for the migration defects in sorbs1-deficent cells (Figure 5F,G).

**Figure 5:**
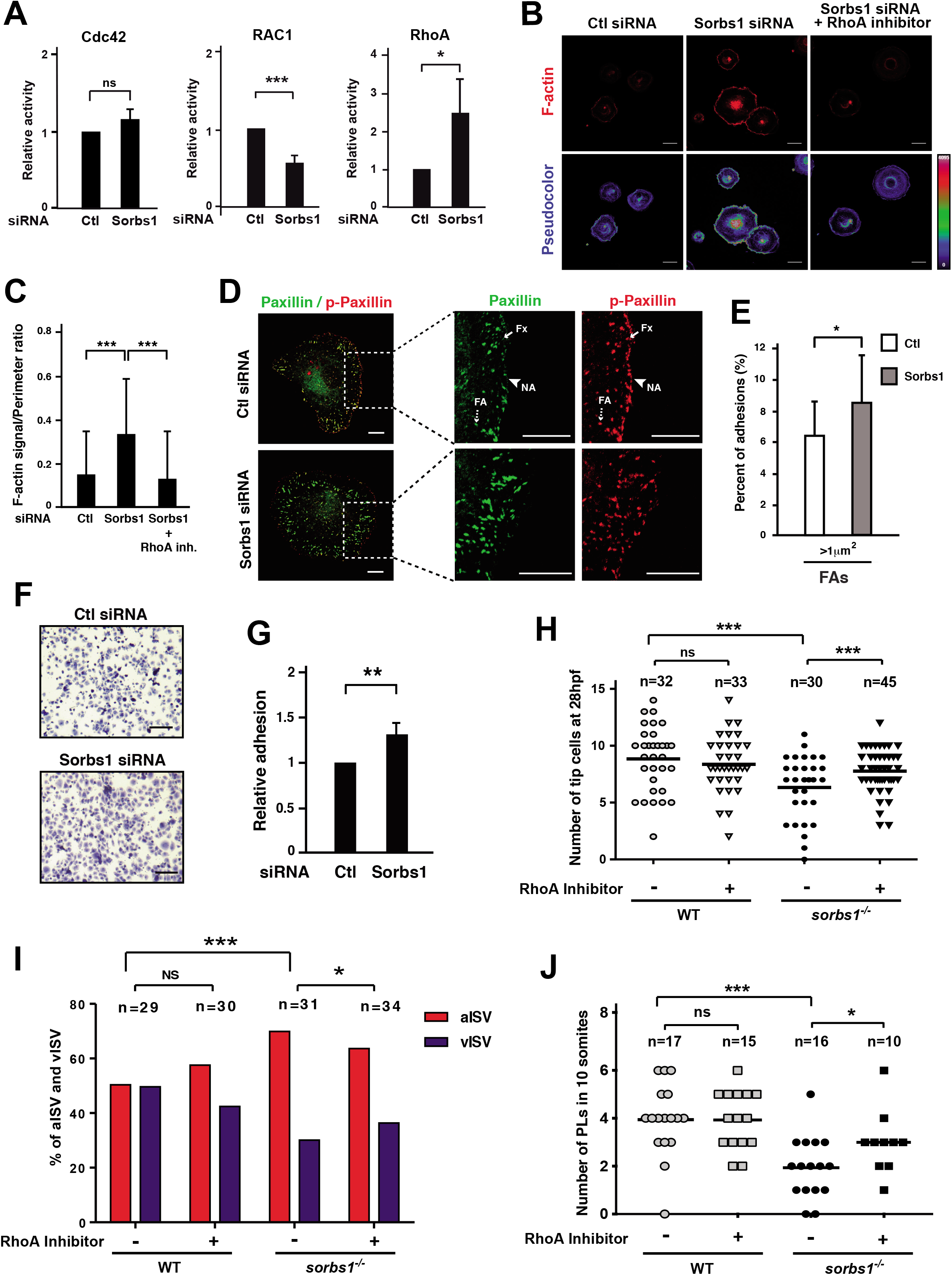
Sorbs1 controls EC adhesive properties via RhoGTPases *in vitro* and *in vivo*. (A) Cdc42, Rac1 and RhoA activity in HUVECs transfected with control (Ctl) or *Sorbs1* siRNA. Histogram is from Western blot densitometric analysis of three independent experiments and represent the ratio between bound active- and total amount of each RhoGTPase in the lysate, relative to control cells. (*** *P<0.001, *P<0.05*, ns=non-significant, Student’s t test). (B) Confocal pictures of peripheral F-Actin (phalloidin staining) in Ctl or Sorbs1 siRNA transfected HUVECS treated (+) or not (-) with the C3 RhoA inhibitor. Images are shown using an intensity-based color look-up table (bottom, from blue = low to red = high). Scale bars represent 25 μM. (C) Quantification of the relative peripheral F-actin signal calculated based on images acquired as in (B) (n=48, 51 and 31 cells; *** *P< 0.001*, Student’s t test). (D-E) Adhesion complexes were analyzed by confocal microscopy (D) after immunostaining of paxillin and phosphor-paxillin in HUVECs transfected with control or *Sorbs1* siRNA. Scale bars are 10 μm. Nascent adhesions (NA) and focal complexes (Fx) are identified (solid arrows) by their small size, peripheral location and high p-Paxillin/Paxillin ratio content. Larger and more mature focal adhesion (FA, dashed arrows) were defined as bigger than 1 μm^2^ and their proportion in each condition was quantified (E). (n=21; * *P< 0.05*, Student’s t test). (F-G) Representative micrographs (F) and quantification (G) of adhesion assays performed with HUVECs transfected with control siRNA or with siRNA against *Sorbs1* as described in the method section. Scale bars represent 100 μm. (n=3 independent experiments; *\*\*P<0.01*, Student’s t test). (H,I) WT and *sorbs1^-/-^* embryos were treated (+) or not (-) with RhoA inhibitor at 26 (H) or 28 (I) hpf and the number of sprouting cells at the edge of developing CVP at 28 hpf (H) or the percentage of aISV/vISV at 48 hpf (I) were quantified (n=number of embryos; *\*\*\*P<0.001*, * *P<0.05;* ns=non-significant; Mann–Whitney U-test (H); χ2 with Yates correction(I)). (J) Quantification of the number of PLs in 10 somites at 54hpf in WT and *sorbs1^-/-^* embryos injected with RhoA inihibitor or left untreated (n=number of embryos; *\*\*\*P<0.001*, * *P<0.05;* ns=non-significant; Mann–Whitney U-test).

Expression of Sorbs1 was induced during cell adhesion onto fibronectin (Supplementary Figure S5K). This process triggers formation of the FAK-Src complex^56^, which is known to induce activation of Rac1 and transient suppression of RhoA, thus promoting adhesion disassembly at cell protrusions. As Sorbs1 localizes to FAs and interacts with FAK, Src and several of their substrates at ligand-bound integrin adhesions^35,39,48,57^ we assessed the activity of this complex upon Sorbs1 depletion. We found that activation of FAK, Src and their downstream target ERK was decreased in Sorbs1-depleted cells (Supplementary Figure S5L).

Altogether these data suggest that Sorbs1 participates in the FAK-Src signaling module, which controls the balance between RhoA and Rac1 activities and regulate adhesion dynamics during EC migration. In that case, one should expect that preventing hyperactivation of RhoA would rescue the defects associated with Sorbs1 deficiency. To test this hypothesis *in vivo*, we treated zebrafish embryos at 26 hpf with the C3 transferase RhoA inhibitor and examined the formation of the vascular structures originating from EC sprouting from the PCV. We used low doses of the C3 RhoA inhibitor, which had no significant impact on the vascular development of wild-type embryos (Figure 5H-J). In contrast, treatment with C3 significantly improved the number of sprouting ECs in the developing CVP and the proportion of vISVs in *sorbs1* mutants (Figure 5H,I). Similarly, RhoA inhibitor injection improved lymphangiogenesis in *sorbs1^-/-^* embryos as PL formation was significantly increased (Figure 5J). In agreement with our hypothesis, these observations altogether demonstrate that the PCV sprouting defects associated with *sorbs1* deficiency are in part mediated by RhoA hyperactivation.

## Discussion

Although previous studies have described inactivation of SoHo family members in various animal models^58–61^ the authors did not specifically examine blood vessels and no references were made to a potential vascular phenotype. Here, using the zebrafish model, we provide the first line of *in vivo* evidence that Sorbs1 is crucial for developmental angiogenesis and lymphangiogenesis in vertebrates. We show that *sorbs1* mutant embryos exhibit specific defects in the lymphatic and venous trunk networks that correlate with edema development and impact larvae survival. The endothelial function of Sorbs1 appeared to be cell autonomous and conserved throughout vertebrates as the defects in the zebrafish vasculature could be rescued by endothelial re-expression of the human ortholog. During our investigations we found enrichment of Sorbs1 in the zebrafish endothelial compartment. However, Sorbs1 is a ubiquitous protein that interacts with multiple promiscuous cytoskeleton components and is present in actin-based structures of several cell types^35,39^. Complementary to the endothelium, we do not exclude that additional cellular compartments might be affected in *sorbs1* knock-out embryos. Further studies would therefore be needed to better understand how these proteins might be regulated in a cell type- and/or physiological environment-specific manner.

The vertebrate vasculature is established through temporally and spatially defined angiogenic waves, during which new angiogenic structures grow out from pre-existing vessels. These new vessels do not always maintain the lymphatic, venous or arterial identity of the parental vessel. In that aspect, the process of trunk secondary sprouting from the PCV is particularly illustrative. Indeed, neighboring ECs within the PCV sprout simultaneously towards the dorsal plate. However, these sprouts rapidly diverge and develop into two differently fated structures: the intermediate pool of midline PLs that later give rise to the trunk lymphatic system and the vessel connections between the PCV and the ISVs that establish the venous intersomitic network. Induction of endothelial Prox1a expression (and therefore presumably transcriptional re-programing) in some ECs from the PCV precedes secondary sprouting and correlates with their lymphatic fate^7,9^. Yet, recent data suggest that this specification could rather rely, at least partially, on a Notch-driven heterogeneity preexisting in the primary aISVs that are approached by the venous sprouts: aISV-forming ECs differ in their polarity and mobility before secondary sprouting, impacting on ISV/PCV/DA connection outcome^18^. In contrast, high temporal resolution imaging revealed that secondary sprouts of both fates display very similar behavior early during migration, with most lymphatic sprouts connecting transiently to the aISVs before assembling into PLs^18^. In agreement with the idea that egression of lymphatic and venous sprouts from the PCV share common signaling pathways and downstream intracellular mechanisms, failure in PL formation is often associated with impaired arterio-venous ISV patterning ^22^,^27,31,32,45,62^. Our data show that lack of Sorbs1 has no obvious effect on Prox1a specification. However, it severely impairs secondary sprouting capacities of ECs from both venous and lymphatic fates, which would position Sorbs1 as part of the cellular machineries required for the early morphogenetic events underlying EC secondary sprouting from the PCV.

Migration of these secondary lympho-venous sprouts has been shown to rely on Vegfc signaling^22,27,47^. Even though this master lymphangiogenic driver also controls the earlier process of Prox1 a induction^22,63^, some Vegfc downstream effectors impact LECs behavior without affecting Prox1a specification^30,63^, suggesting that Vegfc can use distinct routes to instruct different lymphangiogenic steps. Sorbs1 appears to regulate lymphangiogenesis independently of Vegfc signaling. Indeed, *sorbs1^-/-^* mutant embryos remained highly responsive to ectopic Vegfc induction. Whereas this result does not exclude that Sorbs1 could be involved in endogenous Vegfc signaling during lymphangiogenesis, genetic interaction experiments demonstrated that Sorbs1 and Vegfc very likely act in distinct pathways. The dichotomy between Sorbs1 and Vegfc signaling is rather unique as other known regulators of PLs and TD formation appear to mostly function within the Vegfc pathway.

Examination of other bed specific angiogenic processes gave us additional insights about potential signaling pathways in which Sorbs1 could function during blood and lymphatic vessel formation. Although we did not perform systematic analysis, head vascularization, which displays strong organotypic signatures including during facial lymphangiogenesis^8,64^, appeared unaffected in *sorbs1^-/-^* mutants, with the absence of periorbital edema^65^. In contrast, we observed defects in the development of the CVP and the SIVP networks, two structures originating from ventral migration of ECs out of the axial vein. Since initiation of CVP formation^19^ and SIVP outgrowth^20,21^, both of which are affected in Sorbs1 mutants/morphants, specifically rely on BMP signaling, we tested and observed a clear involvement of Sorbs1 in BMP-induced ectopic sprouting from the CVP. Does that imply that the defects related to sprouting events from the PCV exhibited by *sorbs1^-/-^* embryos, including lympho-venous dorsal migration, are BMP-dependent? Photo-conversion experiments have revealed that the same population of progenitor cells located within the ventral side of the PCV that generates lymphatic PLs also migrates rostrally to be incorporated into the SIVP^9^. Although PL and SIVP formations occur in opposite direction, suggesting distinct cues and signaling pathways, BMP-related transcriptional activity has been described in lymphatic sprouts budding from the cardinal vein during mouse embryonic development^66^. Apart from a morpholino-based study suggesting a role for type II BMP receptors in PL formation^67^, the potential involvement of BMP signaling during the early steps of lymphatic network formation has never been precisely characterized beyond LEC specification^68,69^. In the light of our findings, it would be interesting to investigate this possibility and test the role of BMP signaling in the lymphatic defects observed in *sorbs1* mutant embryos.

The context-specific phenotypes in the vasculature of *sorbs1* mutants are highly remarkable and suggest that this cytoskeleton-associated protein participates in establishing endothelial cell specificities required throughout PCV-derived secondary venous and lymphatic beds. ECs might use different cellular and molecular mechanisms to establish organ-specific vasculature. For instance, extension of filopodia is crucial to the EC “sheet-like” migration during CVP morphogenesis but is dispensable for the “phalanx-like” migration during ISV development^70,71^. Lympho-venous sprouting and CVP formation are particularly sensitive to microtubule cytoskeleton-associated polarity^72^. Disconnection of the leading ECs from the original vessel during SIVP formation is not observed during formation of ISV and CVP and could suggest distinct mechanisms^73^. Using cell culture experiments, we showed that *Sorbs1^KD^* ECs display altered adhesion dynamics and migration. Interestingly, the CVP phenotype in *sorbs1* mutants is strikingly similar to that of zebrafish embryos lacking various components of the ECM such as fibrillins^74,75^. Additionally, zebrafish mutant for Polydom/svep1, a large protein involved in cell adhesion to the extracellular matrix, failed to form PL and TD lymphatic structures due to sprouting impairment of properly specified LECs^32,76^. This raises the intriguing possibility that venous plexus and lymphatic network formation might be particularly sensitive to alterations of integrin-mediated cell-ECM adhesions.

Our study not only provides an important cellular and developmental *in vivo* context for cytoskeleton regulation by Sorbs1, but it also discloses underlying mechanistic aspects. Indeed, we demonstrate that Sorbs1 acts upstream of RhoGTPases to control EC actomyosin cytoskeleton and migratory behavior. Prior to this work, only few studies had alluded to potential connections between SoHo proteins and RhoGTPases signaling^50,51,77,78^. Although extension of filopodia and migration of leading ECs during BMP-induced CVP morphogenesis was shown to be dependent on the Cdc42 RhoGTPase^71^, we found that Cdc42 activity was not affected in the absence of Sorbs1 in ECs. In contrast, we show that Sorbs1 controls the RhoA-Rac1 balance, through the FAK-Src pathway. Integrin-mediated activation of the FAK-Src complex during cell spreading and migration stimulates Rac1 activity and maturation of focal complexes into stable adhesions^79^. Consistent with the idea that it participates in FAK-Src activation, Sorbs1 protein levels are highly and transiently induced following integrin engagement onto fibronectin. FAK and Src also control phosphorylation of p190RhoGAP and in addition to Rac1 activation, Sorbs1 might affect focal adhesion turnover through repression of RhoA activity^80,81^. Together with the well-described antagonistic regulation of Rac1 and RhoA, these findings would be consistent with the reciprocal increase in RhoA and decrease in Rac1 activities that we observed in Sorbs1-depleted ECs. On this particular matter, it is noteworthy that we were unable to correlate the higher RhoA activity following depletion of Sorbs1 with increased stress fiber formation, cellular contractility or overall ROCK1/MLCK activity. Instead, we observed a rather spatially restricted effect, with Sorbs1-depleted cells exhibiting denser peripheral bundles of actin filaments. Interestingly, asynchronous activation of Rac1 and RhoA activities at the cell edge is essential for membrane protrusions formation and motility of non-endothelial cells^82^. Based on our observations it is tempting to speculate that Sorbs1 might contribute to the spatial coordination of RhoA and Rac1 activities within migrating ECs during vascular network expansion. In the absence of Sorbs1, local increase in RhoA and decrease in Rac1 activities would be expected to reduce lamellipodia dynamics and membrane protrusive activity, thus impairing EC migration. Importantly, we show that defects in PCV secondary dorsal and ventral sprouting associated with Sorbs1 knockout can be partially rescued by a RhoA inhibitor, indicating that RhoA activation is causal in the vascular phenotype of *sorbs1^-/-^* embryos and suggesting that Sorbs1 affects common endothelial cell properties during these processes. How RhoA regulation by Sorbs1 is integrated in the signaling pathways governing these processes is still an open question but it is worth noting that while Erk activation has been described to participate in LEC migration^47^ and CVP formation^19^, adhesion-triggered phosphorylation of Erk is reduced in *Sorbs1^KD^* ECs.

In summary, the results reported here indicate that the Sorbs1 protein participates in key molecular pathways driving stage- or context-specific regulation of EC morphogenic properties during vascular development. More specifically, we identify Sorbs1 as a novel genetic regulator of developmental lymphangiogenesis that functions independently of Vegfc signaling. Better understanding of these pathways and identification of novel actors provide new opportunities that can be exploited for vascular normalization strategies in various diseases.

## Materials and Methods

### Cell Culture and transfection

Human Umbilical Vein Endothelial Cells (HUVECs), Human Dermal Microvascular Endothelial Cells (HDMECs), Human Mammary Epithelial Cells (HMECs), Human Umbilical Artery Endothelial Cells (HUAEC), Human embryonic kidney 293 cells (HEK 293), Human Dermal Lymphatic Microvascular Endothelial Cells (HMVEC-dLyAd) and HeLa cells were obtained from Lonza. All functional assays were performed with HUVECS which were grown at 37°C in endothelial basal medium (EBM) supplemented with hydrocortisone (1 μg/ml), bovine brain extract (12 μg/ml), gentamicin (50 μg/ml), amphotericin B (50 ng/ml) epidermal growth factor (10 ng/ml) (Lonza) and 10% Fetal Bovine serum (FBS, Perbio). Transfections of siRNA were performed using the GeneTrans II (MoBiTec) reagent according to the manufacturer’s protocols.

Except for scratch-wound assays, all functional assays were performed on fibronectin-plated HUVECs. Briefly, cells were harvested, left 30 min in suspension to recover from trypsinisation and seeded onto fibronectin-coated dishes for 30 min.

### Migration assays

Migration assays were performed as described [ref 30]. Briefly, for the scratch assay a confluent HUVEC monolayer was wounded 48hrs after siRNA transfection, using a sterile P200 tip to create a cell-free zone. For each wound, two different fields were photographed just after injury (t = 0 h) and 16 h later. Quantification of cell migration was made by measuring the percentage of area recovery using ImageJ software in 12 fields from 3 independent experiments.

### Adhesion assay

Adhesion assays were performed essentially as described [ref 30] with slight modifications. Forty-eight hours after transfection, HUVECs were seeded on fibronectin precoated-wells for 30 min. After extensive washing with PBS, remaining cells were stained with cristal violet. The dye was released by cell permeabilization and directly proportional to the number of cells, dye concentration was measured by reading absorbance at 560nm.

### Proliferation assays

Forty-eight hours post transfection with siRNA, a colorimetric MTS assay was performed on HUVECs following the manufacturer’s protocol (CellTiter 96 AQ ueous One Solution Cell Proliferation Assay, Promega) in 96 wells. Alternatively, semi-automatic cell counting assessment of proliferation was performed. Briefly, 24h post-transfection with siRNA, cells were seeded at 20000 cells/well in 24-well plates in triplicate. Cells were counted using the Scepter 2.0 Handheld Automated Cell Counter (Millipore) over a 2-day period and proliferation curves were generated by plotting the average cell number over time.

### Immunohistochemistry

AccuMax Array (A301 VI) slides were stained with goat anti–human Sorbs1 (Abcam, ab4551) antibody. Slides were incubated overnight in optimized dilutions of primary antibodies in Antibody Diluent (Dako, S2022). Peroxidase-conjugated anti–goat Ig (Vector) was then added for 1 hour. Revelation was performed using diamino-3, 3’ benzidine (DAB) according to standard protocols. Images were acquired by using a FSX100 microscope (Olympus).

### Immunofluorescence

For immunofluorescence experiments, HUVECs were seeded onto fibronectin coated coverslips 48 h after siRNA transfection. Cells were fixed after 30 min in 4% paraformaldehyde, permeabilized with PBS-TritonX 0.1% and incubated overnight with the appropriate primary antibody dilutions in PBS-BSA 4%. Cells were then incubated with appropriate secondary antibody dilutions for 1 h. After washing, cells were mounted with Mowiol (Sigma) and processed for immunofluorescence using a confocal Nikon A1R.

### Zebrafish

Adult fish and embryos were carried on according to EU regulations on laboratory animals. All animal experiments were approved by the animal welfare committee of the University of Liège and the Université libre de Bruxelles (ULB). The zebrafish lines used in this study were: *Tg(fli1a:eGFP*)^*y1*83^, *Tg(hsp70l:bmp2b)^84^, TgBAC(prox1a:KalTA4-4xUAS-ADV.E1b:TagRFP*)^*nim*85,86^, vegfchu5055^29^.

### Generation of knockout lines using CRISPR7Cas9 system

Cas9 mRNA and guide RNAs (gRNAs) were synthesized as described in Jao et al.^87^. Briefly, the Cas9 mRNA was synthesized by *in vitro* transcription using the T3 mMESSAGEmMACHINE Kit (#AM1348, Ambion). The primers for the generation of DNA templates of gRNAs were designed through the CHOPCHOP software, and a T7 promoter sequence was added to the 5’-upstream of the gRNA sequence. The gRNA was digested by BamHI and then submitted to in vitro transcription using MEGAshortscriptT7 kit (#AM1354, Ambion). The size and quality of the capped mRNA and gRNA were confirmed by electrophoresis through a 2% (w/v) agarose gel. After this, 300ng/μl of Cas9 mRNA and 100 ng/μl of gRNA were co-injected into one cell-stage zebrafish embryos. Embryos were derived from the transgenic line *Tg(fli1a:eGFP)y1* cross. The injected embryos were raised to adulthood. To test mutagenesis efficiency we genotyped the zebrafish by extracting the DNA from their fin (FIN-CLIP), followed by PCR and heteroduplex melting annealing (HMA) gel. F0 fish were crossed with *Tg(fli1a:eGFP)y1* fish to generate heterozygous F1 progeny, which were then genotyped by HMA gel and DNA sequencing. Heterozygous F1 zebrafish were crossed with the aim to generate homozygous mutant fish.

### Morpholino injection

One-cell stage *Tg(fli1a:eGFP)y1* embryos were injected with 5 ng of Sorbs1 splice-blocking (5’-TCCCCAAATGCTCTTCTTACCAGTA-3’) and control morpholino (5’-CCTCTTACCTCAGTTACAATTTATA-3’). We performed rescue experiments by injecting RNA molecules (60ng/μl) from *in vitro* transcription reactions using linearized PCS2+ vector coding for human Sorbs1.

### RNA extraction and PCR amplification

RNA was extracted from zebrafish embryos using Trizol reagent (Invitrogen) according to the manufacturer’s protocol. RNA from HUVECs was prepared using the nucleospin RNA kit (Macherey Nagel). RNA integrity and concentration were assessed by spectrophotometry analysis (Nanodrop, Thermo Scientific). Reverse transcription reactions were done using the RevertAid H Minus First Strand cDNA Synthesis Kit (Fermentas) with random hexamer primers. The cDNA was then submitted to quantitative real time PCR using Sybrgreen technology (Eurogentec) on a Stepone apparatus (Applied Biosystems) or to end-point PCR amplification followed by gel electrophoresis analysis.

Primers used for end-point PCR are: Zebrafish *sorbs1:* ATCATCGATGTGCACTAACGTG (Forward) and CTCCAGCAGAGGGCACAG (Reverse). Primers used for quantitative real-time PCR are: Zebrafish sorbs1: GCCAGGAAAGTCTTCAGTGC (Forward) and TCTGCTTCACCGTCACTCAC (Reverse); Zebrafish *prox1a:* TGTCATTTGCGCTCGCGCTG (Forward) and ACCGCAACCCGAAGACAGTG (Reverse). Zebrafish *elfa*: CTTCTCAGGCTGACTGTGC (Forward) and CCGCTAGCATTACCCTCC (Reverse). Primers used for mutagenesis efficiency analysis by PCR are: TGAGACTCCAGCAGACATGG (Forward) and ACAATTACAGCTGGAGAACTACA (Reverse).

### Whole mount in situ hybridization

An antisense RNA DIG-probe was generated by transcription from linearized pCS1 vector containing Sorbs1 coding sequence using SP6 RNA polymerase kit where UTPs were labelled with digoxigenin (DIG) (Roche, 11175025910).

Whole mount in situ hybridization was performed in 12, 24 and 48 hpf embryos and in 3dpf larvae. Every time point was fixed with paraformaldehyde 4% overnight at 40C and then dehydrated and rehydrated through methanol and PBS 1x (Gibco)-Tween 5% washes. Embryos were permeabilized with 10μg/mL of proteinase K and then re-fixed with paraformaldehyde 4%. Antisense probe hybridization was performed using 100 ng of sorbs1-DIG-probes hybridization buffer containing 5% dextran sulfate at 65 °C overnight. The use of a DIG alkaline phosphatase-conjugated antibody (Roche, 11093274910, dilution 1/3000), and its substrates BCIP and NBT, enabled the colorimetric detection of sorbs1 transcript. Pictures were taken with an Olympus SZxX10 stereomicroscope.

### Phenotyping

Embryos were anesthetized with Tricaine 0,4% in order to perform phenotypical analysis. Analysis and pictures of overall zebrafish morphology and edemas were performed under a stereomicroscope. Analyses of zebrafish vasculature were performed under a fluorescent stereomicroscope, whereas confocal pictures were taken on live embryos embedded in low melting point agarose (0.8%) on a confocal Nikon A1R. 3D color projections were done using the volume view-slices mode and the volume view-z depth blending functions of NIS-Element A1R1 Software. Lightsheet Zeiss Z1 was used in order to perform time-lapse video of the emerging secondary sprouts at 36 hpf from the trunk vasculature of zebrafish embryos from crossing *TgBAC(prox1a:KalTA4-4xUAS-ADV.E1b:TagRFP)^nim5^* and *sorbs1^-/-^* homozygotous lines which were embedded in low melting point agarose (0.8%) with Tricaine 0,4%.

For rescue experiments with RhoA inhibitor, control, sorbs1 mutants or morphants embryos were incubated with C3 Transferase RhoA inhibitor (#CT04-A, C3ytoskeleton, Inc.) (1μg/mL) at 26 hpf before analyses of CVP structures at 28 hpf and of the proportion of aISV/vISV at 48 hpf. For PL development rescue, RhoA inhibitor (1 μM) was injected in the circulation of wild type and *sorbs1^-/-^* at 28 hpf, and PLs were quantified with a fluorescent stereomicroscope at 54hpf.

For testing the interaction with vegfc, embryos from *sorbs1^+/-^* and *vegfc^-/-^* crosses were submitted to phenotyping before being genotyped by BsaI digestion (vegfc) or HRM (sorbs1). WT and *sorbs1^-/-^* embryos were injected at the one-cell stage with 200 pg/embryo of human VEGFC plasmid (kindly provided by Dr. Schulte-Merker laboratory) before PL and TD quantification.

### Antibodies and RNA interference (RNAi)

Anti-Sorbs1 was obtained from Abcam (#Ab4551). Anti-PAK4 (#3242), PAK2 (#2608), Src (#2123), ERK1/2 (#9102) and phosphorylated Src (#2101S), paxillin (#2541S) and ERK1/2 (#9101) were purchased from Cell signaling. Anti-paxillin (610051), FAK (#ab72140) and its phosphorylated form (# 44-624G) were from Biosciences, Abcam and Invitrogen, respectively. Non-targeting control siRNA and siRNA duplexes targeting Sorbs1 (5’-UUAAGUCCUGAGUGCUCUUC-3’) were synthesized and purchased from Eurogentec.

### Rho GTPase pull down activity assay

SiRNA-treated HUVECs were cultivated for 30 minutes on fibronectin. After harvesting, total cellular active RhoA levels were measured using the Rho Activity Assay (Cytoskeleton Inc., BK036) following the manufacturer’s guidelines. In short, cell lysate (approximately 500μg of total protein) was incubated for 1 hour at 4 °C with GST-Rhotekin beads. Bound activated RhoA was eluted from the beads and analyzed by western blotting using a RhoA antibody.To measure the levels of active Rac1 and Cdc42 were measured using the Rac1 and CDC42 Activity Assay (Cytoskeleton Inc., BK035 and BK034, respectively), cells were lysed in a buffer containing 5 mM DTT, 50 mM Tris pH 7.2, 1% tritonx-100 (10%), 0,5% deoxycholate (20%), 0,1% SDS (20%) and 500 mM NaCl 5M. Extracts were then incubated 1h at 4°C with GST-PAK beads. Bound activated Rac1 and Cdc42 were eluted from the beads and analyzed by western blotting using dedicated antibodies.

### Statistical Analysis

Unless stated otherwise, experiments were performed at least three times independently and graphs represent means +/− standard deviation. Normality tests were performed and when the data were considered normal, statistical analysis were performed by two tailed Student’s t-test or a Pearson’s chi-squared test. Mann-Whitney U and Wilcoxon rank-sum test were used otherwise.

## Acknowledgments and Sources of Funding

We would like to thank members of the GEC laboratory for helpful and stimulating discussions. We also thank the GIGA-Imaging (Sandra Ormenese) and Zebrafish (Hélène Pendeville) facilities for technical support. We also would like to thank to Dr.Schulte-Merker and laboratory members for providing human vegfc coding plasmid and technical support in zebrafish experiments. This research has been funded by the Interuniversity Attraction Poles Program initiated by the Belgian Science Policy Office (IUAP-BELSPO PVI/28 and PVII/13) and was supported by the Belgian National Fund for Scientific Research and Funds from the Université de Liège. A.B., A.V. and T.O. were FRIA Fellows of the Belgian National Fund for Scientific Research. P.C. was a postdoctoral researcher of the FRS.-FNRSThe CMMI is supported by the European Regional Development Fund and the Walloon Region. Work in the B.V. laboratory is supported by the Queen Elisabeth Medical Foundation (Q.E.M.F.), the FRFS-WELBIO (CR-2017S-05R) and the ERC (GoG Ctrl-BBB 865176).

## Non-standard Abbreviations and Acronyms

bp: base-pair
CV: Cardinal Vein
CVP: Caudal Vein Plexus
DA: Dorsal Aorta
DLAV: Dorsal Longitudinal Anastomotic Vessel
DLLV: Dorsal Longitudinal Lymphatic Vessels
dpf: days post-fertilization
EC: Endothelial Cells
ECM: Extracellular Matrix
ELV: Ectopic Longitudinal Vessel
EV: Ectopic Vessel
FA: Focal Adhesions
Fx: Focal Complexes
GAP: GTPase Activating Protein
GFP: Green Fluorescent Protein
gRNA: guide RNA
hpf: hours post-fertilization
ICV: Interconnecting Vessels
ISV: Intersegmental Vessel
Mo: Morpholino
NA: Nascent Adhesions
PCV: Posterior Cardinal Vein
PL: Parachordal Lymphangioblasts
SIV: Subintestinal Vein
SIVP: Subintestinal Venous Plexus
SoHo: Sorbs Homology Domain
TD: Thoracic Duct

## Supplemental Figure Legends

**Supplemental Figure 1:**
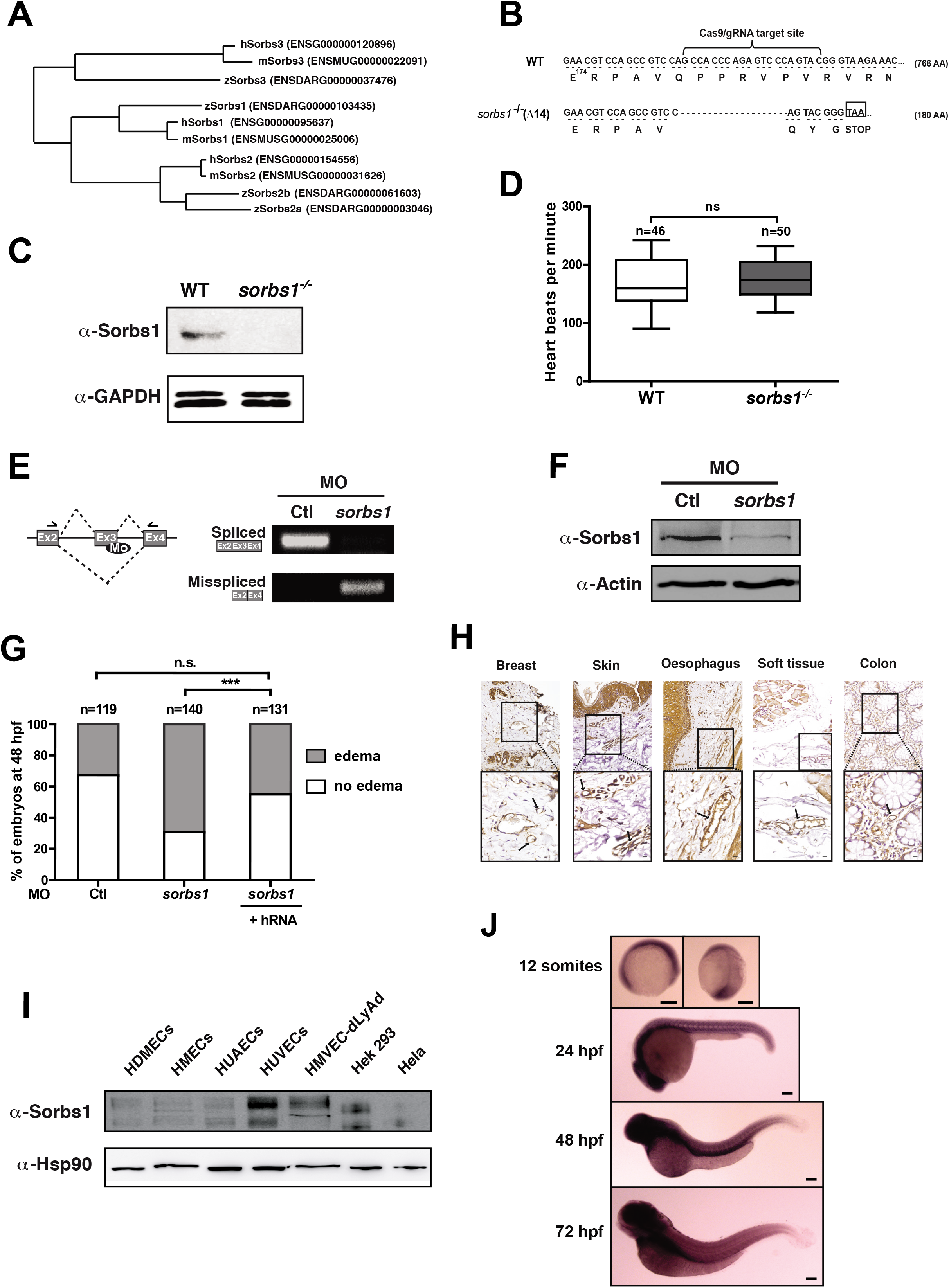
Sorbs1 expression in endothelial cells *in vitro* and *in vivo* and verification of its depletion in the zebrafish model. (A) Phylogenetic tree was constructed from multiple human, mouse and potential zebrafish mRNA of *Sorbs1, Sorbs2* and *Sorbs3* with ClustalW software. One zebrafish ortholog was identified for *sorbs1 (zSorbs1*) and *sorbs3 (zSorbs3*) and two for *sorbs2 (zSorbs2a* and *zSorbs2b).* (B) DNA and amino-acid sequences of the wild-type (WT) and 14-bp deletion (−14) in sorbs1 alleles following CRISPR/ Cas9-based editing. (C) Western blotting analysis of protein extracts from wild-type (WT) and Sorbs1 mutant (*sorbs1^-/-^*) embryos, using anti-Sorbs1 antibody. GAPDH was used as loading control. (D) Quantification of heart beats per minute of wild-type (WT) and *sorbs1^-/-^* homozygote embryos at 2dpf. Depletion of Sorbs1 has no effect on the heartbeat compared to wild-type embryos (n= number of embryos, ns= non-significant, Mann-Whitney U-test). (E) RT-PCR analysis on total RNA from embryos injected with control morpholino (Ctl Mo) or with a splice-blocking morpholino against *sorbs1* (sorbs1 Mo). (F) Western blotting analysis of total protein extracts from 48hpf embryos injected with control (Ctl) and *sorbs1* ATG-blocking Morpholino, using an anti-Sorbs1 antibody. Actin was used as loading control. (G) Quantification of the percentage of edemas-bearing embryos observed at 2 dpf in embryos injected with control or *sorbs1* morpholino, together or not with human Sorbs1 mRNA. (n= number of embryos, *** *P<0.001*, ns= non-significant; Fisher exact test). (H) Expression of Sorbs1 in various human tissues assessed by immunohistochemistry. Typical Sorbs1 staining in endothelial cells is illustrated for the indicated tissues. Boxes correspond to the enlarged area showing expression of Sorbs1 in blood vessels (arrows). Scale bars represent 30 μm and 100 μm respectively in large and zoomed picture. (I) Western blotting analysis of Sorbs1 expression in various human endothelial cells: HDMECs (Human Dermal Microvascular Endothelial Cells), HMECs (Human Mammary Epithelial Cells), HUAECs (Human Umbilical Artery Endothelial Cells), HUVECs (Human Umbilical Endothelial Cells), HMVEC-dLyAd (Human Dermal Lymphatic Microvascular Endothelial Cells), HEK293 (Human Embryonic Kidney 293) and Hela cells. HSP90 was used as a loading control. (J) Whole mount in situ hybridization using a digoxigenin-labeled antisense sorbs1 probe at different time points: 12 somites, 24, 48 and 72 hpf. Scale bars represent 100 μm.

**Supplemental Figure 2:**
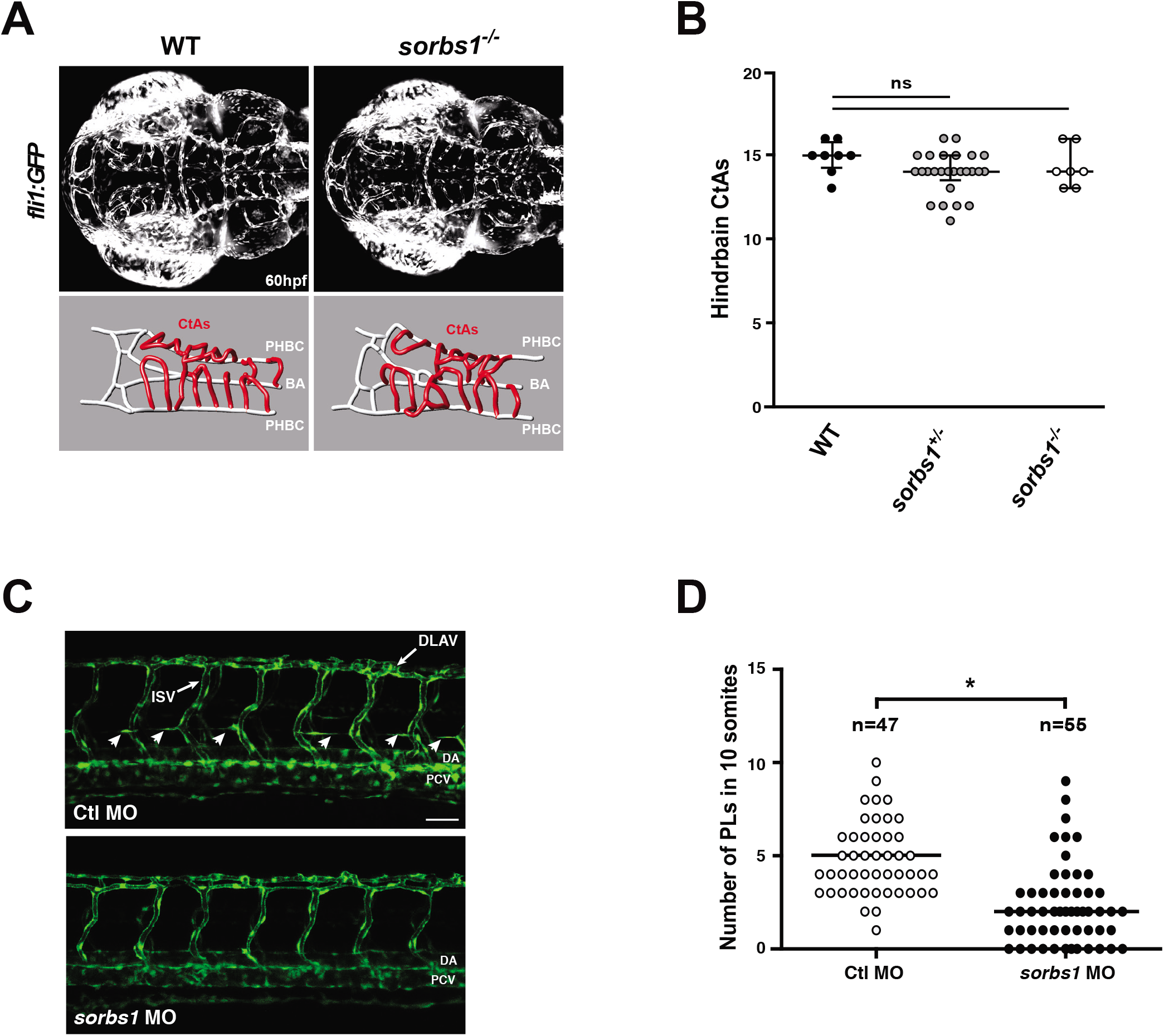
Lack of *Sorbs1* expression does not affect cranial vasculature pattern. (A) Maximal intensity projection of a confocal z-stack of the cranial vasculature of *Tg(fli1a:eGFP)y1* wild-type (WT) and *sorbs1* mutant (*sorbs1^-/-^*) embryos at 60 hpf in dorsal views (anterior to the left) and wire diagram of the brain vasculature in dorso-lateral view. Red vessels in the 3D renderings represent the intra-cerebral central arteries (CtAs) and gray vessels represent the perineural vessels (primordial hindbrain channels: PHBC and basilar artery: BA). (B) Quantification of the corresponding hindbrain CtAs in 8 wild-type (WT), 24 *sorbs1* heterozygous (sorbs1^+/-^) and 7 *sorbs1* homozygous (*sorbs1^-/-^*) embryos at 60 hpf. Error bars represent median ± interquartile range; (n= number of embryos; ns= non-significant, Kruskal–Wallis test). (C) Confocal pictures of the trunk vasculature show no gross vascular defects at 48 hpf in *sorbs1* morphant and Ctl embryos. Defects in PL formation were detected in *sorbs1* morphants (bottom) compared to Ctl embryos (top, white arrowheads). DA: Dorsal Aorta; PCV: Posterior Cardinal Vein; DLAV: Dorsal Longitudinal Anastomic Vessels; ISV: Intersegmental Vessels. Scale bars represent 50 μm. (D) Quantification of PLs was performed at 48 hpf between 10 somites in Ctl and *sorbs1* morphant embryos as imaged in (C) (n=number of embryos; *P<0.05; two-tailed Mann–Whitney *U*-test).

**Supplemental Figure 3:**
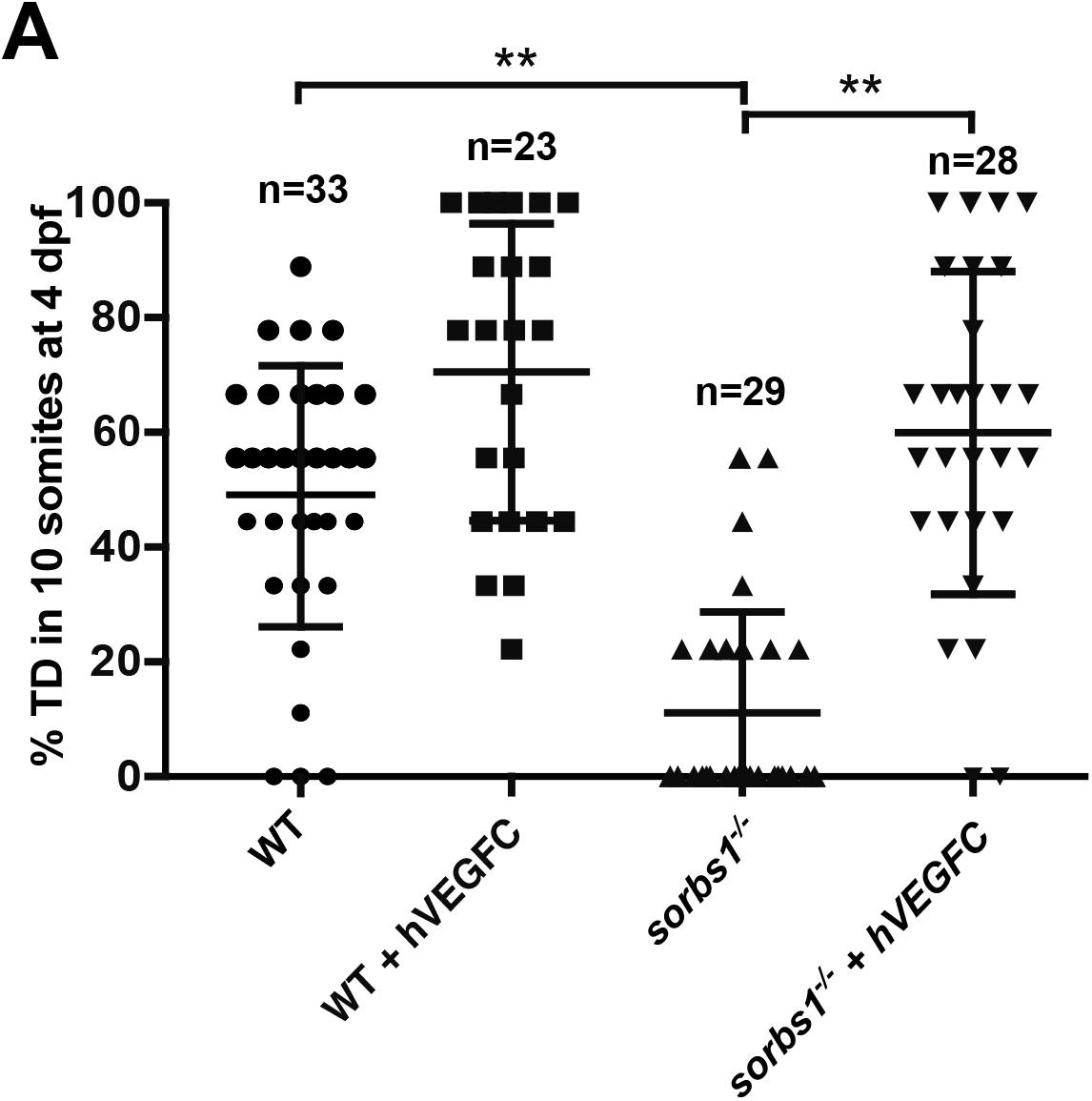
*Sorbs1* depletion does not affect *pro1xa* expression in endothelial cells. (A) Quantification of the trunk TD extent (4dpf) in WT or *sorbs1^-/-^ Tg(fli1a:eGFP)y1* embryos injected or not with human Vegfc. (n= number of embryos; **P<0.01; Mann– Whitney U-test).

**Supplemental Figure 4:**
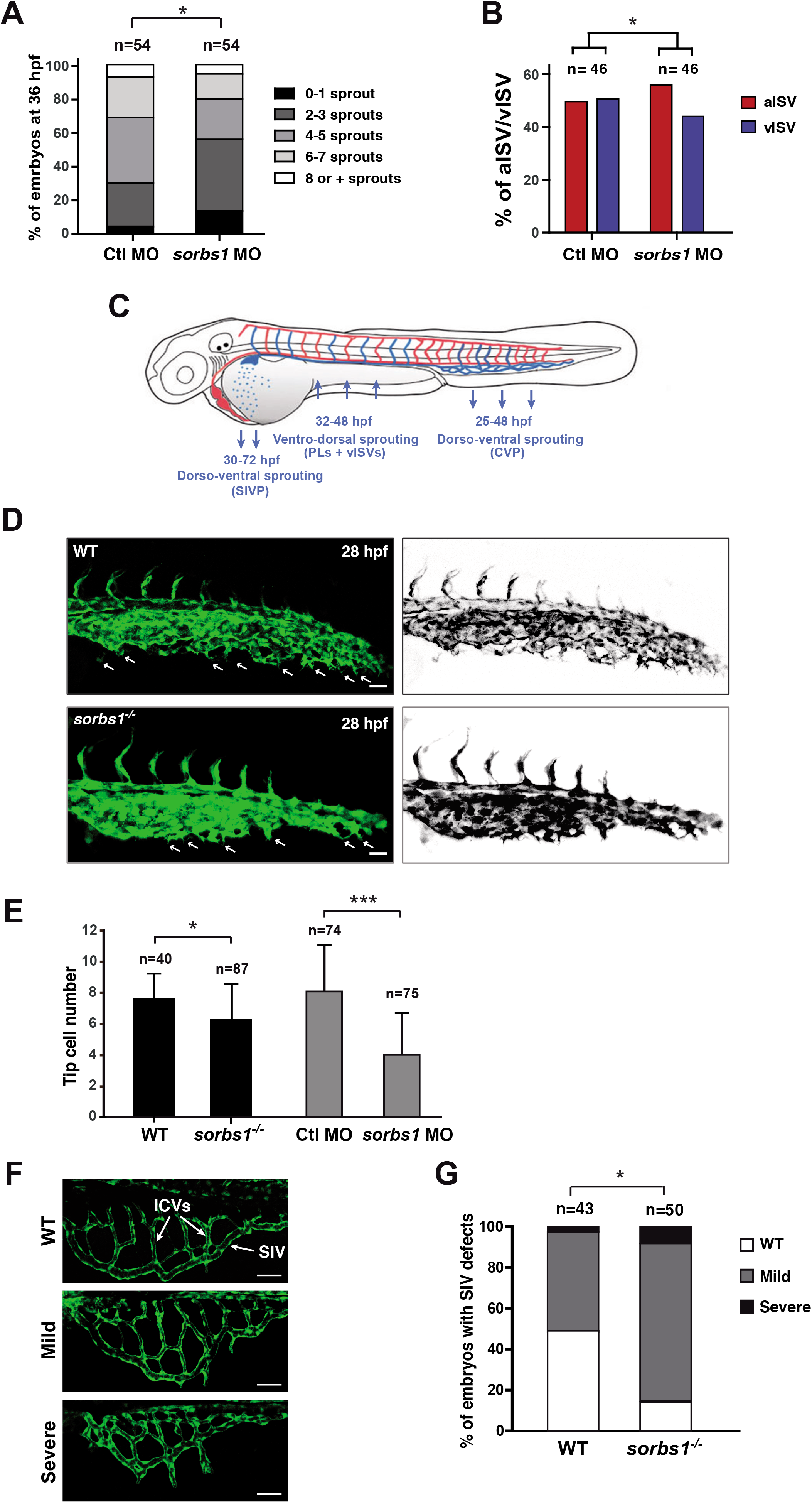
*Sorbs1* depletion leads to defects in secondary sprouting from the PCV. (A) Quantification of secondary sprouts in control and sorbs1-morphant embryos at 36 hpf was established using the five indicated categories. (n = number of embryos; *P<0.05; two-tailed Mann–Whitney U-test). (B) Percentages of vISVs and aISVs were quantified at 48 hpf in a 10 somite region in the trunk of embryos injected with control or *sorbs1* morpholino. (n= number of embryos; *P< 0.01; χ^2^ without Yates correction). (C) Schematic representation of arterial (red) and venous (blue) circulation in the zebrafish embryo. Blue arrows indicate the direction of endothelial cell migration during the formation of PCV-derived angiogenic structures. vISVs: venous Intersegmental vessels; CVP: caudal vein plexus, SIVP: subintestinal venous plexus (D) Confocal imaging of CVP tip cells (white arrows) from 28 hpf wild-type and *sorbs1^-/-^* embryos. Scale bars represent 40 μm. (E) Quantification of tip cell numbers were performed at 28hpf in control and mutant sorbs1 embryos, as well as in embryos injected with control or *sorbs1* Mo (n= number of embryos; * *P< 0.01;* ***P<0.001; two-tailed Mann–Whitney U-test). (F) Confocal pictures of subintestinal plexus of three different phenotypes encountered in WT (first picture) *and sorbs1^-/-^* (second and third picture) *Tg(fli1a:eGFP)y1* embryos taken at 80 hpf. Scale bars represent 40 μm. (G) Quantification of SIV phenotypes as illustrated in F (n=number of embryos, * *P<0.05;* two-tailed Mann–Whitney U-test).

**Supplemental Figure 5:**
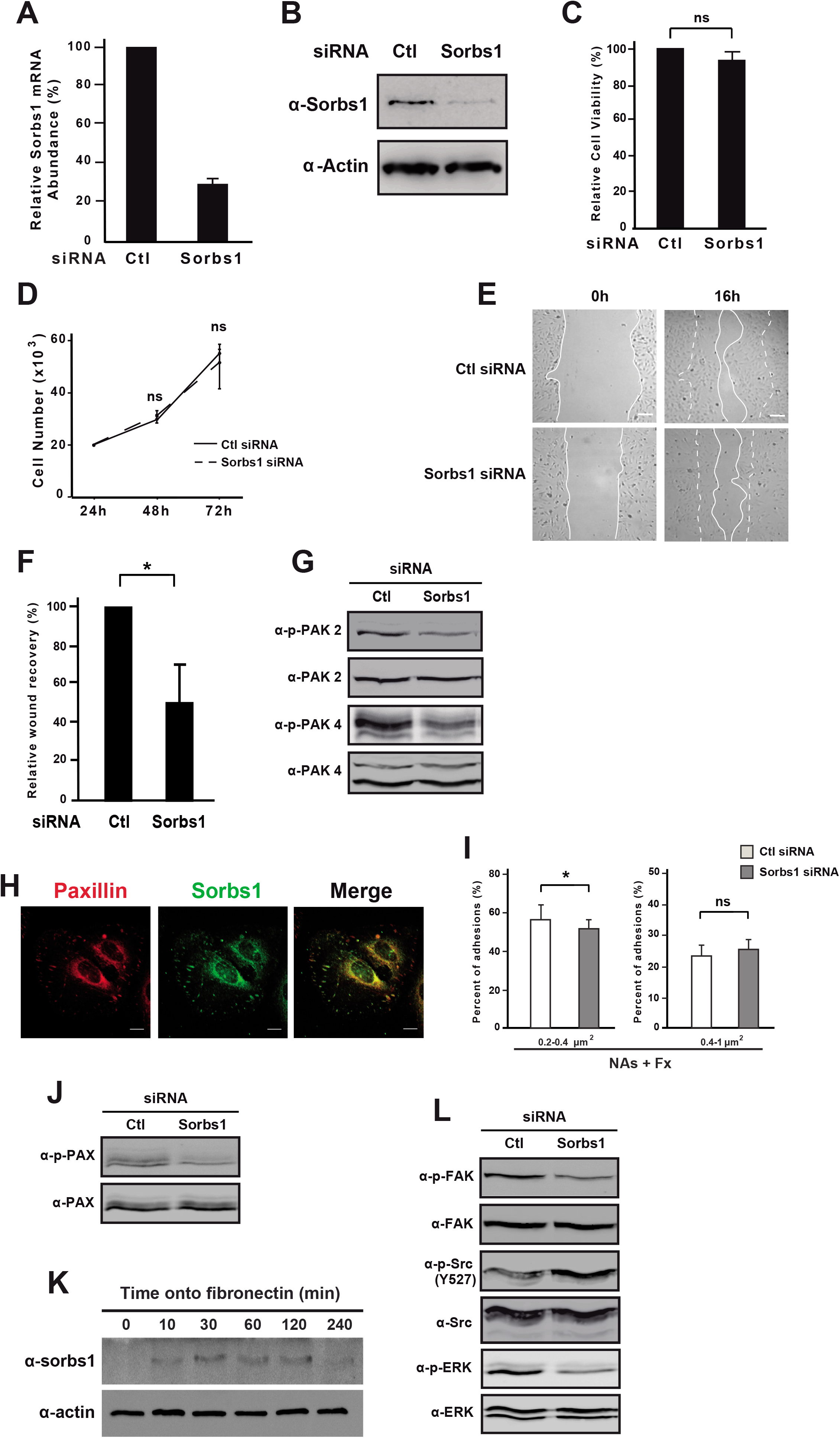
Sorbs1 deletion affects migratory and adhesive properties of ECs. (A) HUVECs were transfected with siRNA targeting *Sorbs1* or with a control siRNA. Efficiency of RNA silencing was assessed by qRT-PCR 48h after transfection. GAPDH was used as internal control. (B) Sorbs1 protein levels were examined in cells described in (A) by Western blotting analysis using Sorbs1 specific antibody. Actin was used as loading control. (C) HUVECs were transfected as in (A). Cell viability relative to control was assessed using an MTS assay, as described in the method section. Results are mean ± SD of 3 independent experiments, each performed in triplicate (ns=non-significant, Student’s t test). (D) HUVECs were transfected as in (A), harvested 24 h after transfection and plated in triplicate at a defined density. Cell number was then assessed by semi-automatic counting (see the method section) at 48 h and 72 h after transfection. Results are presented as the average ± SD increase in cell number, from 3 independent experiments (ns= non-significant, Student’s t test). (E) Micrographs representing HUVECs transfected with control siRNA or with siRNA against *Sorbs1* submitted to a scratch-wound assay. (F) Quantification of the scratch-wound assay as described in E. Histogram represent mean ± sd of 3 independent experiments (*: *P<0.05*, Student’s t test). (G) Rac1 activity in HUVECs transfected with control (Ctl) or *Sorbs1* siRNA. Rac1 activation was assessed by Western blot analysis of PAK2 and PAK4 phosphorylation using phospho-specific antibodies. Total amounts of PAK2 and PAK4 were used as loading controls. (H) Co-localization of Sorbs1 and adhesion complexes was analyzed in HUVECs by confocal microscopy using antibodies specific to paxillin and sorbs1. (I) The size distribution of adhesions was established from HUVECs (n=21) visualized as in Figure 5D. Adhesions were classified into two categories based on their size: [0.2-0.4μm^2^], [0.4-1μm^2^] (NAs + FX). (*: *P< 0.05*, ns= non-significant, Student’s t test). (J) Phosphorylation of Paxillin was assessed in Ctl and Sorbs1-depleted cells using a phospho-specific antibody. Total amount of Paxillin was used as loading control. (K) HUVECs were seeded onto fibronectin for the indicated times and the expression of Sorbs1 was analyzed by Western blotting with a specific antibody. Actin levels were used as loading control. (L) FAK-Src-ERK signaling was assessed in Ctl and Sorbs1-depleted cells. Activated FAK and ERK and inactivated Src were detected using phospho-specific antibodies. Total amounts of the corresponding proteins were used as loading controls.

